# INVA8001, a Novel and Highly Selective Chymase Inhibitor, Ameliorates Liver Inflammation, Fibrosis, and Hyperplasia in *Mdr2* Knockout Mice

**DOI:** 10.1101/2025.11.18.688305

**Authors:** Sameer Sharma, Lixian Chen, Tianhao Zhou, Meenakshi Chawla, Anita Ganjoo, Shunichiro Okada, Salvatore Alesci, Krishnan Nandabalan, Heather Francis

**Author notes:** **Address for correspondence:** Heather Francis, PhD. Author MC was a former consultant to Invea Therapeutics, Inc. Author KN is an employee of Invea Therapeutics and AlphaMeld®Corporation. Authors AG, SO, and SA are employees of Invea Therapeutics. The company holds an exclusive license to ASB17061 (INVA8001) from Daiichi Sankyo Company, Ltd.

## Abstract

**Background and aims:** Mast cells (MCs) play a significant role in autoimmune diseases by mediating inflammatory responses, innate and adaptive responses, angiogenesis, and various pathological processes. MC numbers are significantly increased in chronic liver disease and liver cancer, and degranulation leads to a release of MC mediators, including chymase, which increases inflammation, activates hepatic stellate cells (HSCs), ultimately leading to fibrosis. Primary sclerosing cholangitis (PSC) is a chronic, progressive cholestatic liver disease characterized by inflammation, fibrosis, ductular reaction, and eventual liver failure. MCs and their mediators, particularly chymase, have been implicated in PSC pathogenesis; however, targeting chymase therapeutically has remains largely unexplored. In this study, we aimed to evaluate the efficacy of INVA8001, a highly selective chymase inhibitor, on PSC pathogenesis in the *Mdr2* knockout mouse (*Mdr2^-/-^* mice) model of PSC.

**Approach and Results:** We evaluated the levels of chymase and other MC markers in liver biopsy samples collected from late-stage PSC patients. The effect of chymase inhibition was evaluated using a highly selective and potent small molecule chymase inhibitor, INVA8001, in *Mdr2-/-* mice. Ten-week-old male *Mdr2^-/-^* mice were injected intraperitoneally (IP) with 20 mg/kg of INVA8001 daily for two weeks and varying parameters of disease pathogenesis, including MC activation, inflammation, fibrosis, biliary pathology, and cholestasis were evaluated. In addition, histological, immunohistochemical, biochemical, and molecular analyses were conducted to evaluate the effects of INVA8001 treatment in *Mdr2^-/-^* mice.

The liver biopsies of PSC patients showed an increased number of chymase-positive cells compared with control samples (collected from non-diseased patients). INVA8001 treatment resulted in a reduction of MC accumulation, inflammation, histological damage, fibrosis, ductular reaction, and biliary senescence in *Mdr2^-/-^* mice.

The current data show the pathophysiological role of chymase in PSC and the impact of a selective chymase inhibitor on PSC disease pathogenesis. This can further be extrapolated to various MC-driven diseases such as chronic urticaria, atopic dermatitis, asthma, metabolic dysfunction-associated steatohepatitis (MASH), and eosinophilic gastrointestinal diseases, among others, where chymase is central to the disease pathogenesis.

**Conclusions:** Our findings identify chymase as a key driver of PSC pathogenesis establishing INVA8001 as a promising new therapeutic candidate for hepatobiliary disorders, including PSC. Chymase inhibition simultaneously targets MC activation, inflammation, fibrosis, and biliary senescence, and offers a multifaceted approach to treating PSC and other MC-related disorders.

## Introduction

Primary sclerosing cholangitis (PSC) is a rare, chronic cholestatic disease that gradually progresses from early to end-stage liver disease and detection often occurs at late-stage thus necessitating liver transplantation for patient survival. In the United States, there are currently no Food and Drug Administration (FDA)-approved treatments for PSC management. PSC is primarily characterized by liver inflammation and fibrosis, along with the destruction of the intra- and extrahepatic bile ducts leading to multiple strictures in the biliary tree and eventually cirrhosis. According to 2021 data, the incidence and prevalence of PSC in the general population was 0.75 (95% CI: 0.542–1.05) and 11.16 (95% CI: 7.78–16.02) per 100,000 people, respectively (1). The incidence rate of PSC has significantly increased over time and approximately 40% of patients with this disease ultimately require liver transplants. While PSC is considered a rare disorder, it is associated with high morbidity and mortality, as prognosis is often poor and the condition markedly increases the risk of developing cholangiocarcinoma (2).

Mast cells (MCs) are critical cellular components of the immune system, playing a key role in maintaining tissue homeostasis, by regulating epithelial function and integrity, modulating both innate and adaptive mucosal immunity, and mediating local neuro-immune interactions. Substantial evidence from both human and animal models has validated the role of MCs and their mediators in immune-mediated cholestatic disorders, such as PSC. MCs positive with both tryptase, a MC–specific serine protease and canonical marker of MC activation, and c-KIT, the receptor for stem cell factor (SCF) ligand (bioactive, soluble SCF), required for MC development and survival, infiltrate the fibrotic portal tracts in the livers of PSC patients and their numbers have been positively correlated with disease severity (3). It has been shown that, in the early stages of PSC, MCs predominantly localize around the bile ducts, whereas in the later stages of the disease, MCs are primarily found in the fibrous septa. An increase in MC markers such as c-KIT and the high-affinity IgE receptor, FceR1, and MC mediators such as chymase, tryptase and histamine have been observed in livers from PSC patients, as well as in the genetic murine model of PSC (*Mdr2*^*-/-*^ mice). Furthermore, treatment with cromolyn sodium, a MC stabilizer, in *Mdr2*^*-/-*^ mice attenuates the levels of MC mediators, inflammation, tissue damage, and fibrosis (4).

Upon activation by pathogen-associated molecular patterns (PAMPs) and danger-associated molecular patterns (DAMPs), MCs undergo degranulation and release a variety of bioactive mediators including chymase (5). Chymase, a serine protease, plays a pivotal role in the recruitment of various immune cells, such as eosinophils, neutrophils, macrophages, and lymphocytes and initiates a cascade of events that promotes inflammation, epithelial barrier dysfunction and fibrosis (6). Once chymase is released, it triggers the activation of several key substrates including angiotensin II (Ang-II), matrix metalloproteinases (MMP-1, MMP-9), transforming growth factor beta (TGF-β), collagen I, and chemokines such as interleukin-33 (IL-33) (6). In addition, chymase leads to the further proliferation of MCs by enzymatically converting SCF to its bioactive soluble form, which in turn binds to c-KIT receptors on MCs and supports proliferation, migration, survival, and differentiation of MCs (7). In chronic liver injury, the expression of SCF is induced in biliary epithelial cells that line bile ducts showing periductal fibrosis and inflammation as compared to non-affected bile ducts, suggesting a role for cholangiocytes in MC accumulation and proliferation (8).

Chymase and its substrates are implicated in promoting liver inflammation and fibrosis, and chymase inhibitors have demonstrated therapeutic potential in various animal models of liver diseases. For example, chymase activity is induced in inflamed livers of hamsters fed with a methionine- and choline-deficient (MCD) diet and blocking chymase function inhibits Ang-II and MMP-9 activity, as shown in liver biopsies derived from hamsters fed with MCD diet (9). Additionally, chymase inhibition has been shown to alleviate steatohepatitis, limit infiltration of MCs, decrease expression of pro-inflammatory and pro-fibrotic mediators, and restrict fibrosis in the liver in various steatohepatitis animal models (10). Although elevated levels of chymase have been observed in liver samples from PSC patients, there is no published literature to date examining the effect of chymase inhibition on PSC pathogenesis.

INVA8001 (formerly ASB17061; in-licensed by Invea Therapeutics, Inc. from Daiichi Sankyo Company, Ltd.) is a complex organic molecule with a benzene ring substituted with a side chain containing a butyl group and, 4-diazepan-1-yl ring (a 7-membered ring containing nitrogen) with an attached chloro-methoxybenzyl group (**Figure 1**). The compound is crystallized with acetic acid molecules which are incorporated into its structure as solvent molecules. INVA8001 is an orally administered, highly potent and selective chymase inhibitor with a high safety index. *In vitro* primary pharmacology studies demonstrate that INVA8001 is a potent and highly selective inhibitor of the MC-chymase enzyme and shows remarkable selectivity over related enzymes (**Table 1**) (11).

**Table 1:**
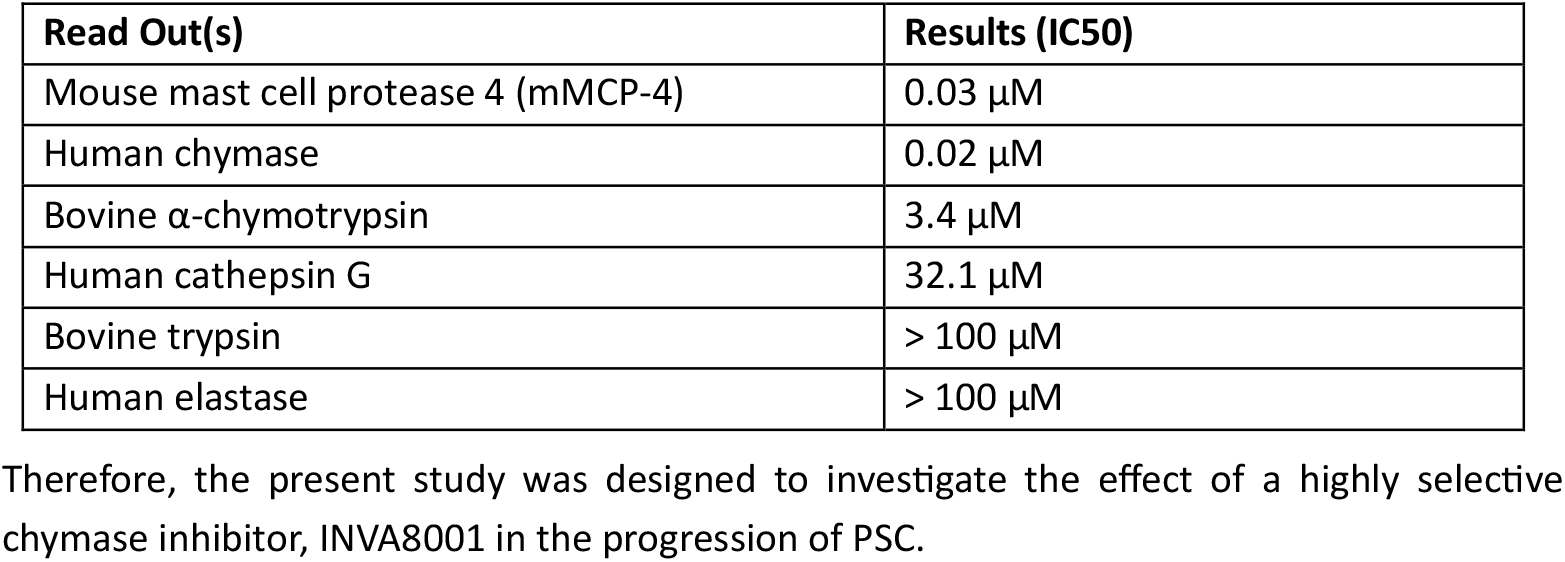
Selectivity of INVA8001.

| Read Out(s) | Results (IC50) |
| --- | --- |
| Mouse mast cell protease 4 (mMCP-4) | 0.03 μM |
| Human chymase | 0.02 μM |
| Bovine α-chymotrypsin | 3.4 μM |
| Human cathepsin G | 32.1 μM |
| Bovine trypsin | > 100 μM |
| Human elastase | > 100 μM |

**Figure 1:**
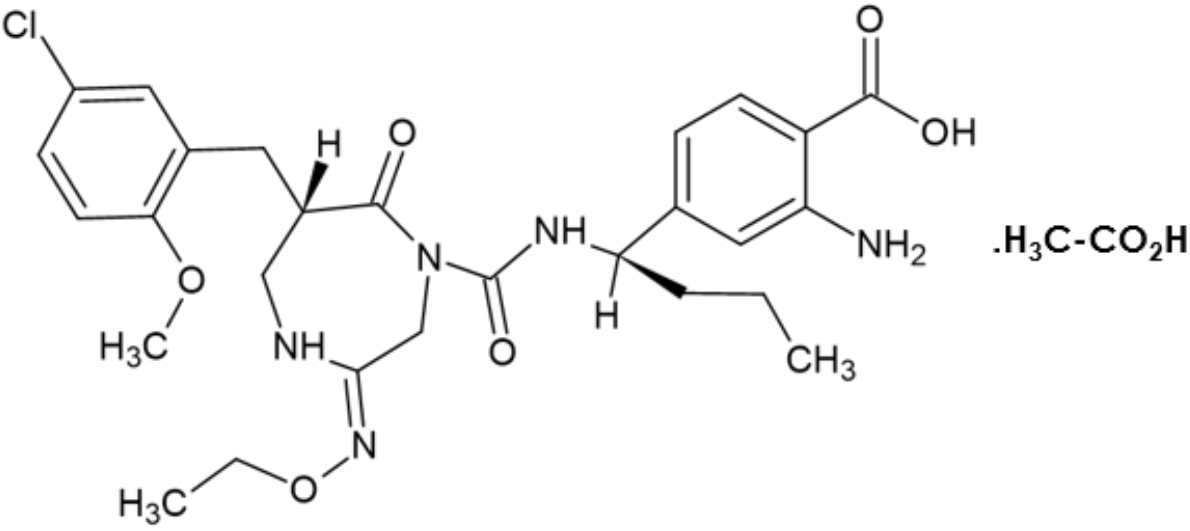
Structure of INVA8001, a novel chymase inhibitor.

## Materials and Methods

### Materials

Study drug, ASB17061 (INVA8001), was supplied as a gift sample by Daiichi Sankyo Company, Ltd. with >95% purity. Reagents were purchased from Sigma-Aldrich (St. Louis, MO) unless otherwise indicated. Information for all antibodies used is described in Supplemental Table 1. The Quick-RNA Miniprep Plus the Kit for RNA isolation was purchased from Zymo Research (Irvine, CA). Selected mouse primers were purchased from Qiagen (Valencia, CA). The iScript cDNA Synthesis Kit (170-8891) and iTaq Universal SYBR Green Supermix were purchased from Bio-Rad (Hercules, CA). The information on real-time PCR (*qPCR*) primers used is listed in the Supplemental Table 2.

### Animal models

Animal procedures were performed according to protocols approved by Indiana University School of Medicine Institutional Animal Care and Use Committee. Ten-week-old male *Mdr2*^*-/-*^ mice (n = 8) and Friend Virus B NIH Jackson (FVB/NJ [FVB/wt control], n = 8) mice were treated with daily IP injections of INVA8001 (formulated in 0.5% methyl cellulose and dosed at 20 mg/kg) for two weeks.

In all groups, serum, liver, cholangiocytes, and cholangiocyte supernatants were collected as described (12), (13). The collected liver was divided into three portions: (i) snap-frozen tissue; (ii) OCT-embedded blocks (5 µm cryosections) for immunofluorescence (IF); and (iii) formalin-fixed, paraffin-embedded (FFPE) blocks (4 µm sections) for immunohistochemistry (IHC).

### Human samples

Human explant liver tissues from both male and female patients were collected from either normal controls (liver n=5) or late-stage PSC patients (liver n=5) under the Institutional Review Board (IRB) approved protocol at Indiana University School of Medicine (IUSM). Informed consent was collected from patients for the human specimens that were utilized in this study. An additional three healthy control liver samples were purchased from Sekisui Xeno Tech (Kansas City, KS). The samples were stained by IHC for chymase and tryptase in FFPE liver sections.

### Evaluation of liver damage, ductular reaction and biliary senescence

Liver damage was assessed by hematoxylin and eosin (H&E) staining in FFPE mouse liver sections and scored in a coded fashion to determine portal inflammation. Serum and liver total bile acids (TBAs) were determined using a TBA Assay Kit (Colorimetric, Sigma-Aldrich). Ductular reaction was evaluated in FFPE mouse liver sections by IHC for CK-19 to specifically mark bile ducts. Stained slides were scanned by a digital scanner (Aperio AT2 scanner; Leica Biosystems). CK-19 positive area was quantified using the Image-Pro Premier software (Rockville, MD). Biliary senescence was evaluated by: (i) IF for cyclin dependent kinase inhibitor 2a (Cdkn2a, gene for p16) expression in frozen liver sections co-stained with CK-19 (to mark bile ducts) and (ii) mRNA expression of cellular senescence markers, (Cdkn2a; gene for p16), cyclin dependent kinase inhibitor 2c (Cdkn2c, gene for p18) and galactosidase, beta 1-like (Glb1l) in isolated mouse cholangiocytes (3) (13).

### Assessment of liver fibrosis

Collagen deposition was determined by Fast Green/Sirius Red staining in FFPE mouse liver sections and quantified by the Image-Pro Premier software (Rockville, MD). Liver fibrosis was also determined by measuring hydroxyproline levels in total liver samples using the hydroxyproline assay kit (MAK008-1KT; Millipore Sigma, Billerica, MA). To detect activated HSCs, IF for desmin (co-stained with CK-19) was performed in frozen liver sections with semi-quantification (14), (15). The mRNA expression of the fibrosis markers, collagen type I alpha 1 (Col1a1) and tissue inhibitor of metalloproteinases 1 (Timp1) and transforming growth factor beta 1 (Tgfb1) was measured in total mouse livers by qPCR.

### Evaluation of liver inflammation and MC infiltration

Liver inflammation was evaluated by IHC for F4/80 in FFPE mouse liver sections and positive cells were quantified by Image-Pro Premier software (Rockville, MD). Liver MC infiltration was detected with IHC for tryptase β-2 and mucosal MC chymase-1 in liver sections. MCs were semi-quantified as tryptase β-2 (Tpsb2) or chymase (mMCP1) positivity cell number/total area with Image-Pro Premier software (Rockville, MD). MC activation was determined by *q*PCR for Fc receptor, IgE, high affinity I, alpha polypeptide (*Fcer1a*), *mMCP1* and *Tpsb2* in total liver samples.

### Statistical analysis

All data are expressed as the mean ± SD. Differences between groups were analyzed by unpaired Student’s *t* test when two groups were analyzed and one-way ANOVA when more than two groups were analyzed, followed by an appropriate *post hoc* test. Statistical analyses and generation of graphs were performed with GraphPad Prism (version 9.2.0; GraphPad Software, LLC; San Diego, CA). The level of significance was set at *P* < 0.05.

## Results

### Chymase-positive MCs are elevated in human PSC liver explants

We first investigated the expression levels of chymase and other MC markers, such as tryptase, in liver biopsy samples derived from explanted PSC patients and healthy controls using IHC. We observed that the number of chymase-positive MCs was significantly higher in PSC patients compared to the controls (**Figure 2A**). Similarly, a marked increase in the number of tryptase-positive MCs were observed in PSC liver samples compared to the healthy controls (**Figure 2B**). Our results corroborate previous studies in PSC patients demonstrating increased expression of MC markers, such as c-KIT, FceR1, chymase, and tryptase (16).

**Figure 2:**
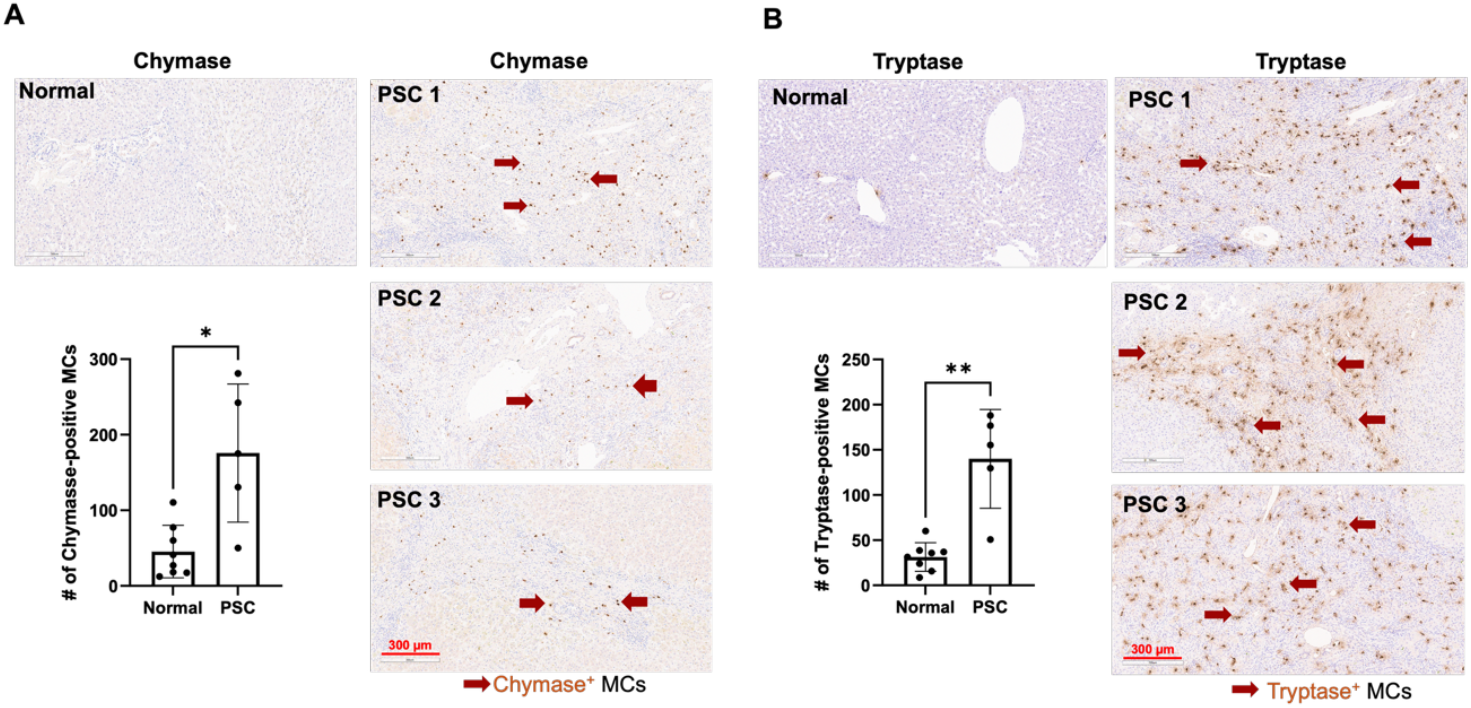
Chymase and tryptase expression in human PSC. (**A**) Representative images for Chymase and (**B**) tryptase IHC staining (×10; scale bar, 300 µm) and semi-quantification (average positive number count for three portal images in per sample) in human samples. Data are mean ± standard deviation. n = 5 human normal control, and n = 8 human PSC. \**P* < 0.05; \*\**P* < 0.01.

### INVA8001 reduces MC accumulation in liver samples from *Mdr2*^*-/-*^ mice

Since we found increased chymase levels in human PSC livers, we evaluated the pharmacological inhibition of chymase in modulating PSC in *Mdr2*^*-/-*^ mice. INVA8001 was administered at a dose of 20 mg/kg (formulated in 0.5% methyl cellulose) for two weeks and injected daily via an intraperitoneal (IP) route of administration (**Figure 3A**).

**Figure 3:**
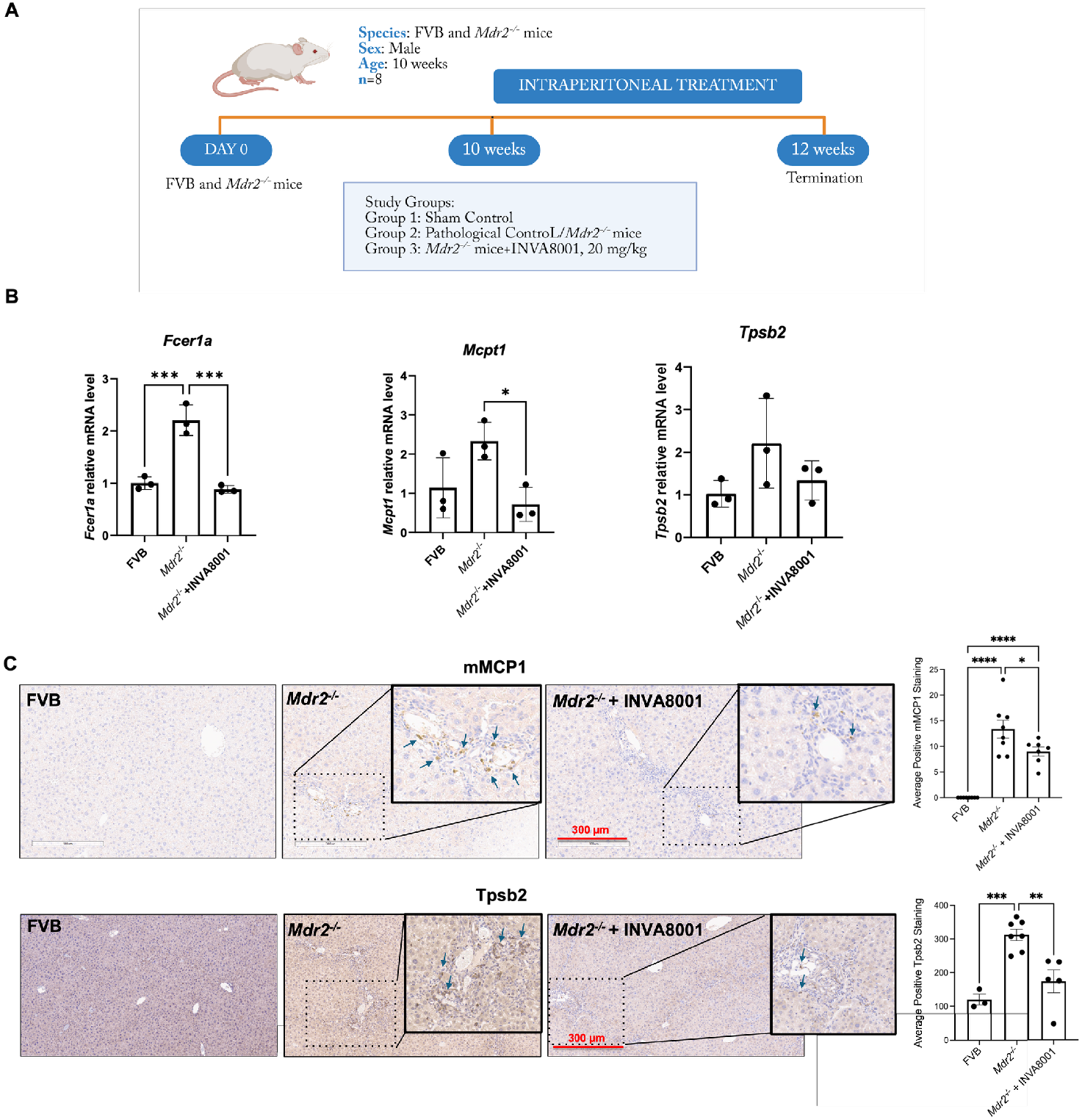
(**A**) Schematic outline of *in vivo* treatment in FVB and *Mdr2*^*-/-*^ mice; (**B**) *Fcer1a, Mcpt1 and Tpsb2* mRNA expression in livers from FVB, *Mdr2*^*-/-*^ and *Mdr2*^*-/-*^ mice treated with INVA8001; (**C**) Representative images for mMCP-1 and Tpsb2 IHC staining (×10; scale bar, 300 µm) and semi-quantification in FVB, *Mdr2*^*-/-*^ and *Mdr2*^*-/-*^ mice treated with INVA8001. Data are mean ± standard deviation. n = 5-8 portal images per sample imaged from n = 3-8 mice for IHC; n = 3 reactions per group in total RNA isolated from livers from n = 8 mice per group. \**P* < 0.05, \*\**P* < 0.01, \*\*\**P* < 0.001, \*\*\*\**P* < 0.0001.

INVA8001 attenuated the expression of *Fcer1a* and MC protease 1 (*Mcpt1)* in the liver samples from *Mdr2*^*-/-*^ mice; however, while the expression of *Tpsb2* decreased, this was not statistically significant (**Figure 3B**). These results suggest that chymase inhibition effectively limits MC accumulation in inflamed livers of *Mdr2*^*-/-*^ mice and may play a very significant role in mitigating MC-driven pathology in PSC.

Further, histological analysis revealed a substantial accumulation of chymase (mMCP1)-positive and tryptase (Tpsb2)-positive cells in *Mdr2*^*-/-*^ mice and INVA8001 dosed at 20 mg/kg, IP led to a significant reduction in the numbers of both chymase-and tryptase-positive cells (**Figure 3C**).

Previous studies have also shown that chymase elicits MC survival, proliferation and activation in part by processing several substrates that function as chemotactic and proangiogenic mediators. Several chymase substrates such as SCF, IL-33, Ang-II, and MMPs promote angiogenesis and these molecules can act as chemotactic mediators (17), (18), (19), (20). For instance, SCF induces MC proliferation and subcutaneous administration of SCF has been shown to enhance MC densities in different organs (7). SCF is also a substrate for chymase and inhibiting chymase prevents cleavage and generation of a mature, bioactive, soluble, form of SCF (18). Additionally, chymase-mediated cleavage of SCF leads to activation of cytokines such as IL-33, which mediates the recruitment of different immune cells including MCs, macrophages, eosinophils and lymphocytes (21).

Together, these findings underscore the multifaceted role of chymase in promoting MC accumulation and activation and support the therapeutic potential of INVA8001 in modulating MC–mediated liver pathology in PSC.

### INVA8001 ameliorates liver damage and inflammation in *Mdr2*^*-/-*^ mice

*Mdr2*^*-/-*^ mice exhibit hallmark histological features of PSC such as lobular damage, necrosis, and inflammation, as revealed by H&E staining. INVA8001 at the tested dose showed an amelioration of histological damage and portal inflammation in *Mdr2*^*-/-*^ mice (**Figure 4A**). Furthermore, TBA levels were shown to be elevated in *Mdr2*^*-/-*^ mice as compared to FVB controls, both in serum and hepatic tissues samples. INVA8001 treatment led to a significant reduction in TBA accumulation at a dose of 20 mg/kg, IP (**Figure 4B**). Although INVA8001 mitigated bile acid accumulation and hepatic damage, no significant changes in liver enzymes were observed (data not shown). An increased number of macrophages were also evaluated by F4/80 staining in the portal tracts of *Mdr2*^*-/-*^ mice and the treatment with INVA8001 significantly reduced the number of infiltrating macrophages (**Figure 4C**).

**Figure 4:**
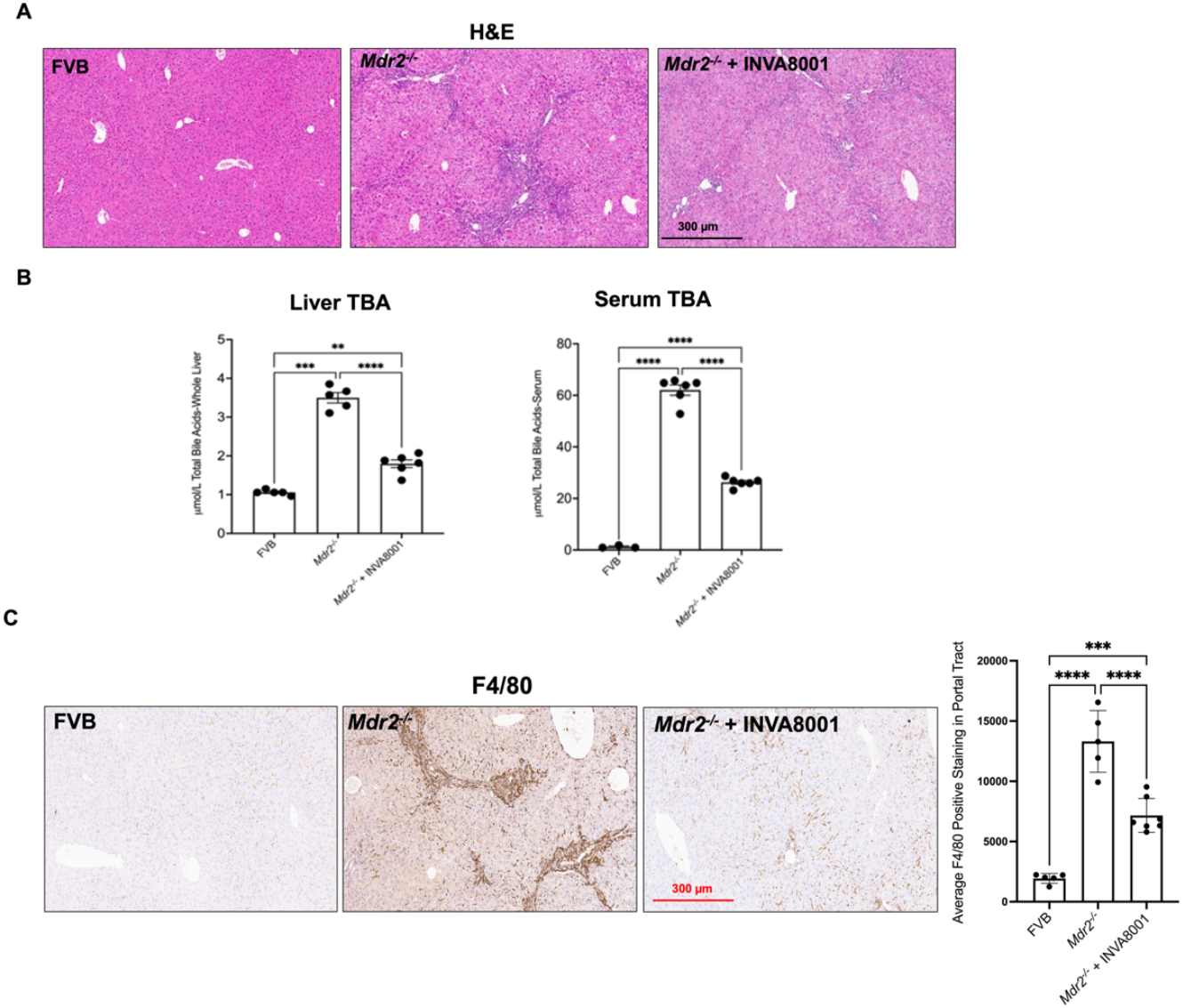
(**A**) Representative H&E staining (×10; scale bar, 300 µm) in mouse livers.; (**B**) liver (left) and serum (right) TBA levels in FVB, *Mdr2*^*-/-*^ and *Mdr2*^*-/-*^ mice treated with INVA8001; (**C**) F4/80 staining (×10; scale bar, 300 µm) and semi-quantification in mouse samples. Data are mean ± standard deviation. n = 5-7 portal areas per mouse imaged from n = 5-7 mice per group for F4/80. \**P* < 0.05, \*\**P* < 0.01, \*\*\**P* < 0.001, \*\*\*\**P* < 0.0001.

Together, these results highlight the detrimental role of chymase in promoting hepatic inflammation and HSC activation. Our findings demonstrate the significant effects of INVA8001 and support its potential as a therapeutic agent to alleviate liver injury and immune cell infiltration in PSC.

### INVA8001 attenuates fibrosis in liver in *Mdr2*^*-/-*^ mice

Next, we assessed the impact of INVA8001 on fibrosis in *Mdr2*^*-/-*^ mice. We evaluated fibrosis by FG/SR staining and quantified hydroxyproline levels (a biochemical marker of collagen deposition), in liver samples. INVA8001 treatment resulted in an amelioration of hepatic fibrosis, as measured by decreased FG/SR staining and lowered hydroxyproline content, compared to untreated mice (**Figure 5A and 5B**). Furthermore, the gene expression levels of key fibrotic markers—*Col1a1, Timp2*, and *Tg@1* in the liver was also significantly reduced following INVA8001 treatment (**Figure 5C**), demonstrating that chymase inhibition prevents hepatic fibrosis in animal models of PSC. These findings corroborate with previous studies that show chymase activates several pro-fibrotic substrates, including collagen and TGF-β, both of which play critical roles in the fibrotic process and chymase inhibition attenuates these effects in varying animal models of steatohepatitis (20).

**Figure 5:**
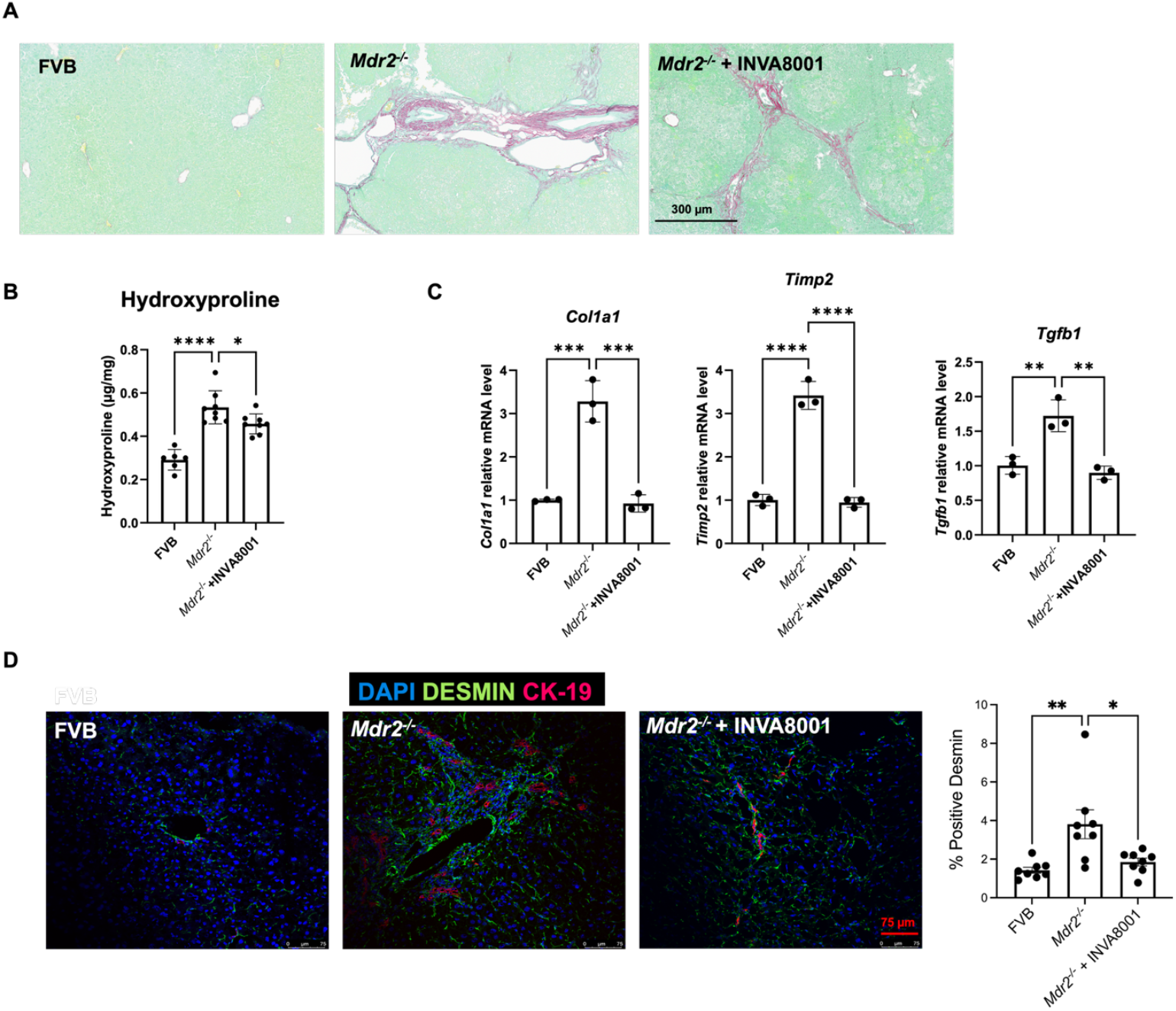
(**A**) Representative FG/SR staining (×10; scale bar, 300 µm) in mouse samples; (**B**) Hydroxyproline content in mouse samples (*μ*g/mg); (**C**) *Col1a1, Timp2* and *Tg91* mRNA expression in total liver from mouse samples and (**D**) Representative IF images (×20; scale bar, 75 µm) for co-staining of CK-19 and desmin in mouse samples and semi-quantification for positive desmin area. Data are mean ± standard deviation. n = 3 reactions per group in total RNA isolated from n = 6 mice; n = 6-8 mice per group for hydroxyproline; n = 3 portal areas per mouse imaged from n = 8 mice per group for desmin. \**P* < 0.05, \*\**P* < 0.01, \*\*\**P* < 0.001, \*\*\*\**P* < 0.0001.

*Mdr2*^*-/-*^ mice are also characterized by the activation of HSCs and portal tract inflammation, both of which contribute to liver fibrosis. Desmin, a marker for HSC activation, showed strong immunoreactivity in *Mdr2*^*-/-*^ mice livers, which was also significantly reduced with INVA8001 at a dose of 20 mg/kg, IP. (**Figure 5D**).

### 2INVA8001 prevents ductular reaction and biliary senescence

Ductular reaction and biliary senescence are both hallmarks of PSC. In PSC, cholangiocyte response to genetic and environmental insults leads to a heterogeneous response with a subpopulation acquiring a proliferative and reactive phenotype, while another subpopulation undergoes senescence and growth arrest, contributing to hepatic inflammation, fibrogenesis and disease progression (22). Here, we evaluated the effect of blocking chymase activity on proliferating ductal and lobular hepatocytes, as well as senescence-associated markers. INVA8001 (20 mg/kg, IP) treatment led to a significant reduction in bile duct mass, as evidenced by a decrease in CK-19-positive bile ducts in *Mdr2*^*-/-*^ mice (**Figure 6A**). Furthermore, INVA8001 administration also reduced the expression of senescence-associated genes, such as *Cdkn2a, Cdkn2c, Tg@1* and *Glb1l* in isolated cholangiocytes from *Mdr2*^*-/-*^ mice (**Figure 6B**).

**Figure 6:**
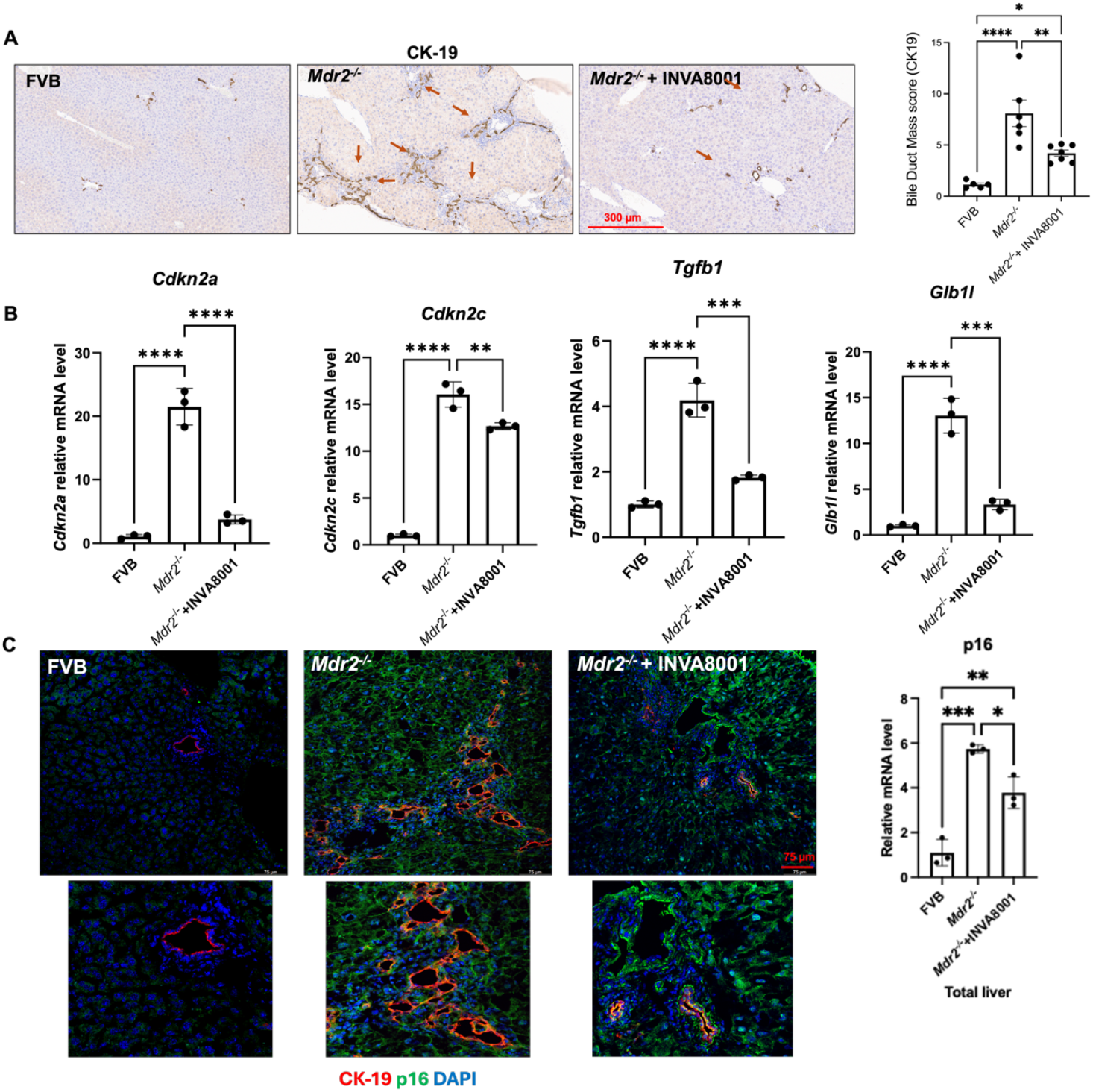
(**A**) Staining (×10; scale bar, 300 µm) and semi-quantification of CK-19 in mouse samples; (**B**) *q*PCR for *Cdkn2a, Cdn2c, Glb1l* and Tgfb1 in isolated cholangiocytes from mouse samples; (**C**) Co-staining (×20; scale bar, 75 µm) for p16 and CK-19 in mouse samples and (**D**) *q*PCR for p16 in total liver from mouse samples. Data are mean ± standard deviation. n = 4–5 portal areas per mouse imaged from n = 5-8 mice per group for CK-19; n = 3 reactions per group in total RNA isolated from isolated cholangiocytes from n = 8 mice per group; n = 3–5 portal areas imaged from n = 3–4 mice per group for p16/CK-19; \**P* < 0.05, \*\**P* < 0.01, \*\*\**P* < 0.001, \*\*\*\**P* < 0.0001.

p16 (encoded by *Cdkn2a*) is a cyclin-dependent kinase inhibitor that regulates and slows down cell cycle progression. INVA8001 treatment also limits the level of p16 immunoreactivity in the livers of *Mdr2*^*-/-*^ mice at a dose of 20 mg/kg, IP (**Figure 6C**), representing a viable strategy to target these pathological processes in PSC.

## Discussion

PSC is a rare, chronic, progressive cholestatic liver disease characterized by increased MC infiltration, inflammation, fibrosis, destruction of both intra- and extra-hepatic bile ducts, and eventual liver failure. Despite its clinical significance and increasing prevalence, effective pharmacological therapies for PSC remain lacking.

Chymase, an enzyme produced and released by MCs, plays a central role in the pathogenesis and progression of hepatobiliary diseases, including liver fibrosis, MASH, and PSC. Elevated hepatic chymase levels correlate with fibrosis severity, and the presence of chymase-positive MCs in fibrotic areas, indicates their direct involvement in disease progression (10) (23). Functionally, chymase exerts both paracrine and autocrine effects, influencing inflammation, extracellular matrix remodeling, and local tissue signalling through its proteolytic activity. Although most effects are paracrine, autocrine loops further amplify MC activation and tissue injury (6) (24) (25).

In the present study, we evaluated the chymase pathway—mediated by MCs—as a critical regulator of PSC pathogenesis and demonstrated that pharmacological inhibition of chymase using a selective small-molecule chymase inhibitor, INVA8001, confers significant therapeutic benefit in a murine model of PSC. Our findings reveal that chymase levels are significantly elevated in liver biopsies from PSC patients compared to healthy controls. These results are consistent with prior studies indicating increased infiltration of chymase- and tryptase-positive MCs in the fibrotic portal tracts of PSC livers. Thus chymase activity substantially increases upon MC activation and activates TGF-β1 (26). INVA8001 effectively suppresses this response, suggesting broader therapeutic effect when compared to the MC stabilization (cromolyn sodium) effects in PSC-induced damage (16).

Using *Mdr2*^*-/-*^ mice, we demonstrated that INVA8001 limits MC accumulation in the liver. INVA8001 decreased the number of chymase- and tryptase-positive cells, and suppressed expression of MC markers such as *FcεRI, Mcpt1*, and *Tpsb2*. These findings are mechanistically supported by prior reports that chymase regulates MC proliferation and survival possibly through substrates like SCF and IL-33 (27) (28) (29) (30). By interrupting this feed-forward loop, chymase inhibition may disrupt the chronic inflammatory environment that characterizes PSC.

Importantly, INVA8001 ameliorated histological liver injury in *Mdr2*^*-/-*^ mice, as evidenced by reduced lobular damage, necrosis, and portal inflammation. While liver enzyme levels were unchanged, INVA8001 treatment led to significant reductions in serum and hepatic bile acid levels, suggesting improvement in bile acid homeostasis. This was accompanied by decreased activation of HSCs, as shown by diminished desmin staining, and a marked reduction in macrophage infiltration, indicating broad immunomodulatory effects of chymase inhibition on the hepatic microenvironment.

One of the most clinically relevant observations was the ability of INVA8001 to attenuate hepatic fibrosis, a hallmark characteristic of PSC progression. Chymase inhibition by INVA8001 reduced collagen deposition (measured by FG/SR staining and hydroxyproline content) and downregulated fibrogenic genes including *Col1a1, Timp2*, and *Tg@1*. These effects are consistent with chymase’s known ability to activate pro-fibrotic mediators such as collagen and TGF-β and align with previous reports of chymase inhibitors reducing fibrosis in models of steatohepatitis.

Furthermore, INVA8001 had a profound impact on ductular reaction and biliary senescence, two emerging contributors to PSC pathology (29) (31). The reduction in CK19-positive ductal mass and the suppression of senescence-associated genes (*Cdkn2a, Cdkn2c, Glb1l, Tg@1*) highlight a novel role for chymase in driving cholangiocyte dysfunction. Given that cellular senescence contributes to chronic inflammation and fibrosis, these findings position chymase inhibition as a promising strategy to interrupt the cycle of cholangiocyte injury and maladaptive repair.

Chymase, a serine protease cleaves and activates various substrates, including MMP-9, Ang-II, IL-33, TGF-β, and collagen I. MMP-9 mediates the infiltration of neutrophils, macrophages, and T cells, facilitating inflammatory responses (32) (33), and Ang-II regulates the maturation and activation of macrophages, dendritic cells, and lymphocytes (34). IL-33 is responsible for promoting infiltration and activation of mast cells, basophils, innate lymphoid cells, macrophages, lymphocytes, and eosinophils in allergic diseases (21). In addition, chymase also enhances eosinophil viability, infiltration, and activation (35) and the infiltration of immune cells combined with the activation of TGF-β and collagen I, which ultimately drives fibrosis (6).

Moreover, chymase is known to cleave SCF into its, mature, bioactive, soluble form, which binds to c-KIT receptors on MCs leading to proliferation, migration, survival, and differentiation (36). By inhibiting chymase, INVA8001 is expected to reduce the recruitment of damaging immune cells—particularly MCs and macrophages—diminish chronic inflammation and epithelial barrier damage, and alleviate cholangiocyte activation, proliferation, and senescence. This, in turn, promotes tissue remodeling and reduces the likelihood of fibrosis across PSC and potentially other related diseases.

Collectively, our data support the conclusion that chymase plays a central, multifactorial role in the progression of PSC, affecting MC activation, inflammation, fibrosis, ductular reaction, and biliary senescence. INVA8001, a selective and a potent chymase inhibitor, demonstrated consistent efficacy across various pathological features in *Mdr2*^*-/-*^ mice establishing a strong preclinical rationale for further development of chymase inhibitors as therapeutic agents in PSC and potentially other mast cell related diseases.

PSC is an orphan disease with no approved medical treatment and the current standard of care for these patients falls into three categories: medical, endoscopic, and surgical. Medical treatment is focused on the off-label use of ursodiol (ursodeoxycholic acid), which at high doses, has been shown to worsen the clinical outcome of the disease, including making some patients ineligible for transplant in combination with drugs aimed at providing symptomatic relief (e.g., cholestyramine, antihistamines, rifampicin, opioid antagonists, and bisphosphonates). Endoscopic therapy is primarily used to manage dominant strictures, and a liver transplant (surgical) is the only therapeutic option for patients for enhanced survival (37) (38) (39) (40) (41). Moreover, given the costs and limited organ availability from donor programs, it is reserved for those with end-stage liver disease only. Thus, there is a high unmet need for safe and effective medical options, which are disease-modifying, halt disease progression to cirrhosis, and delay or avoid liver transplant.

INVA8001 offers a safe and effective oral small molecule therapy compared to the existing therapies for PSC. It is a well-tolerated drug which has been previously tested in a Phase 2 clinical trial registered at ClinicalTrials.gov (42), for atopic dermatitis, with no significant adverse effects. INVA8001’s favorable safety profile and wide therapeutic window support longer-term evaluation and exploration of potential synergies with approved or investigational anti-fibrotic agents. Additionally, the translational potential of INVA8001 could be expanded through biomarker-guided clinical trials in PSC patients, particularly those exhibiting MC– dominant or fibrotic phenotypes such as chronic urticaria, atopic dermatitis, asthma, MASH and eosinophilic gastrointestinal diseases, among others, where chymase is central to the pathogenesis of the disease. In conclusion, INVA8001 can be an effective therapeutic strategy to alleviate MC-mediated disease pathologies and further studies are warranted.

## Abbreviations

Ang-II: Angiotensin II
Cdkn2a: Cyclin-dependent kinase inhibitor 2a
Cdkn2c: Cyclin-dependent kinase inhibitor 2C
CK-19: Cytokeratin-19
c-KIT: Proto-oncogene receptor tyrosine kinase
CI: Confidence interval
Col1a1: Collagen type I alpha
DAMPs: Danger-associated molecular patterns
Glb1l: Galactosidase, beta 1-like
Fcer1a: Fc receptor, IgE, high affinity I, alpha polypeptide
FFPE: Formalin-fixed, paraffin-embedded
H&E: Hematoxylin and eosin
HSCs: Hepatic stellate cells
IF: Immunofluorescence
IHC: Immunohistochemistry
IL-33: Interleukin-33
ILCs: Innate lymphoid cells
MC: Mast cell
Mdr2: Multidrug resistance transporter 2/ATP binding cassette subfamily b member 4
mMCP1: Mucosal mast cell protease 1
MMP-1: Matrix metalloproteinase-1
MMP-9: Matrix metalloproteinase-9
MASH: Metabolic dysfunction-associated steatohepatitis
PAMPs: Pathogen-associated molecular patterns
PBC: Primary biliary cholangitis
PSC: Primary sclerosing cholangitis
qPCR: Quantitative polymerase chain reaction
SCF: Stem cell factor
TBA: Total Bile Acids
TGF-β: Transforming growth factor beta
Timp1: Tissue inhibitor of metalloproteinase 1
Timp2: Tissue inhibitor of metalloproteinase 2
Tpsb2: Tryptase β-2
WT: Wildtype

## Acknowledgements

Dr. Burcin Ekser, MD, PhD (Professor of Surgery, Surgical Director of Liver Transplant at Loyola University Chicago; Loyola University Chicago Stritch School of Medicine).

---

**Supplemental Table 1.** List of antibodies.

| Antibodies | Application | Source | Catalog no. |
| --- | --- | --- | --- |
| CK-19 | IF | Developmental Studies Hybridoma Bank (Iowa City, IA) | TROMA-III |
| CK-19 | IHC | Abcam | ab52625 |
| Chymase (human) | IHC | Abcam | ab111239 |
| mMCP-1(mouse) | IHC | Santa Cruz | Sc-17041 |
| Tryptase (human) | IHC | Cell Marque Antibodies | 342M-15 |
| Tryptase (mouse) | IHC | LifeSpan BioSciences | LS-B15255 |
| Desmin | IF | R&D Systems | AF3844 |
| F4/80 | IHC | Cell Signaling | 70076S |
| p16 | IF | Abcam | ab189034 |

**Supplemental Table 2.** List of real-time PCR primers.

| Gene | Species | Detected transcript | Source |
| --- | --- | --- | --- |
| *Col1a1* | Mouse | NM_007742 | QIAGEN |
| *Cdkn2a* | Mouse | NM_010233 | QIAGEN |
| *Cdkn1a* | Mouse | NM_008084 | QIAGEN |
| *Cdkn2c* | Mouse | NM_007671 | QIAGEN |
| *Fn1* | Mouse | NM_010233 | QIAGEN |
| *Fcer1a* | Mouse | NM_010184 | QIAGEN |
| *Gapdh* | Mouse | NM_008084 | QIAGEN |
| *Glb1l* | Mouse | NM_029010 | QIAGEN |
| *Mcpt1* | Mouse | NM_008570 | QIAGEN |
| *Tgfb1* | Mouse | NM_011577 | QIAGEN |
| *Timp2* | Mouse | NM_011594 | QIAGEN |
| *Tpsb2* | Mouse | NM_010781 | QIAGEN |

## References

1. Cooper J, Markovinovic A, Coward S, Herauf M, Shaheen AA, Swain M, et al. Incidence and Prevalence of Primary Sclerosing Cholangitis: A Meta-analysis of Population-based Studies. Inflamm Bowel Dis. 2024;30(11):2019–26.

2. Lazaridis KN, LaRusso NF. Primary Sclerosing Cholangitis. N Engl J Med. 2016;375(25):2501–2.

3. Ishii M, Iwai M, Harada Y, Morikawa T, Okanoue T, Kishikawa T, et al. A role of mast cells for hepatic fibrosis in primary sclerosing cholangitis. Hepatol Res. 2005;31(3):127–31.

4. Tsuneyama K, Saito K, Ruebner BH, Konishi I, Nakanuma Y, Gershwin ME. Immunological similarities between primary sclerosing cholangitis and chronic sclerosing sialadenitis: report of the overlapping of these two autoimmune diseases. Dig Dis Sci. 2000;45(2):366–72.

5. Huber M, Cato ACB, Ainooson GK, Freichel M, Tsvilovskyy V, Jessberger R, et al. Regulation of the pleiotropic effects of tissue-resident mast cells. J Allergy Clin Immunol. 2019;144(4S):S31-S45.

6. Dell’Italia LJ, Collawn JF, Ferrario CM. Multifunctional Role of Chymase in Acute and Chronic Tissue Injury and Remodeling. Circ Res. 2018;122(2):319–36.

7. Iemura A, Tsai M, Ando A, Wershil BK, Galli SJ. The c-kit ligand, stem cell factor, promotes mast cell survival by suppressing apoptosis. Am J Pathol. 1994;144(2):321–8.

8. Tsuneyama K, Kono N, Yamashiro M, Kouda W, Sabit A, Sasaki M, et al. Aberrant expression of stem cell factor on biliary epithelial cells and peribiliary infiltration of c-kit-expressing mast cells in hepatolithiasis and primary sclerosing cholangitis: a possible contribution to bile duct fibrosis. J Pathol. 1999;189(4):609–14.

9. Tashiro K, Takai S, Jin D, Yamamoto H, Komeda K, Hayashi M, et al. Chymase inhibitor prevents the nonalcoholic steatohepatitis in hamsters fed a methionine- and choline-deficient diet. Hepatol Res. 2010;40(5):514–23.

10. Takai S, Jin D. Chymase Inhibitor as a Novel Therapeutic Agent for Non-alcoholic Steatohepatitis. Front Pharmacol. 2018;9:144.

11. Tomimori Y, Manno A, Tanaka T, Futamura-Takahashi J, Muto T, Nagahira K. ASB17061, a novel chymase inhibitor, prevented the development of angiotensin II-induced abdominal aortic aneurysm in apolipoprotein E-deficient mice. Eur J Pharmacol. 2019;856:172403.

12. Ishii M, Vroman B, LaRusso NF. Isolation and morphologic characterization of bile duct epithelial cells from normal rat liver. Gastroenterology. 1989;97(5):1236–47.

13. Glaser S, Lam IP, Franchitto A, Gaudio E, Onori P, Chow BK, et al. Knockout of secretin receptor reduces large cholangiocyte hyperplasia in mice with extrahepatic cholestasis induced by bile duct ligation. Hepatology. 2010;52(1):204–14.

14. Owen T, Carpino G, Chen L, Kundu D, Wills P, Ekser B, et al. Endothelin Receptor-A Inhibition Decreases Ductular Reaction, Liver Fibrosis, and Angiogenesis in a Model of Cholangitis. Cell Mol Gastroenterol Hepatol. 2023;16(4):513–40.

15. Zhou T, Meadows V, Kundu D, Kyritsi K, Owen T, Ceci L, et al. Mast cells selectively target large cholangiocytes during biliary injury via H2HR-mediated cAMP/pERK1/2 signaling. Hepatol Commun. 2022;6(10):2715–31.

16. Jones H, Hargrove L, Kennedy L, Meng F, Graf-Eaton A, Owens J, et al. Inhibition of mast cell-secreted histamine decreases biliary proliferation and fibrosis in primary sclerosing cholangitis Mdr2(-/-) mice. Hepatology. 2016;64(4):1202–16.

17. Lefrancais E, Duval A, Mirey E, Roga S, Espinosa E, Cayrol C, et al. Central domain of IL-33 is cleaved by mast cell proteases for potent activation of group-2 innate lymphoid cells. Proc Natl Acad Sci U S A. 2014;111(43):15502–7.

18. Watanabe N, Tomimori Y, Terakawa M, Ishiwata K, Wada A, Muto T, et al. Oral administration of chymase inhibitor improves dermatitis in NC/Nga mice. J Invest Dermatol. 2007;127(4):971–3.

19. Caughey GH, Raymond WW, Wolters PJ. Angiotensin II generation by mast cell alpha- and beta-chymases. Biochim Biophys Acta. 2000;1480(1-2):245-57.

20. Takai S, Jin D. Pathophysiological Role of Chymase-Activated Matrix Metalloproteinase-9. Biomedicines. 2022;10(10).

21. Chan BCL, Lam CWK, Tam LS, Wong CK. IL33: Roles in Allergic Inflammation and Therapeutic Perspectives. Front Immunol. 2019;10:364.

22. Guicciardi ME, Trussoni CE, LaRusso NF, Gores GJ. The Spectrum of Reactive Cholangiocytes in Primary Sclerosing Cholangitis. Hepatology. 2020;71(2):741–8.

23. Takai S, Jin D. Chymase as a Possible Therapeutic Target for Amelioration of Non-Alcoholic Steatohepatitis. Int J Mol Sci. 2020;21(20).

24. Dai H, Korthuis RJ. Mast Cell Proteases and Inflammation. Drug Discov Today Dis Models. 2011;8(1):47–55.

25. Zhao XO, Lampinen M, Rollman O, Sommerhoff CP, Paivandy A, Pejler G. Mast cell chymase affects the functional properties of primary human airway fibroblasts: Implications for asthma. J Allergy Clin Immunol. 2022;149(2):718–27.

26. Chen H, Xu Y, Yang G, Zhang Q, Huang X, Yu L, et al. Mast cell chymase promotes hypertrophic scar fibroblast proliferation and collagen synthesis by activating TGF-beta1/Smads signaling pathway. Exp Ther Med. 2017;14(5):4438–42.

27. Molfetta R, Lecce M, Milito ND, Putro E, Pietropaolo G, Marangio C, et al. SCF and IL-33 regulate mouse mast cell phenotypic and functional plasticity supporting a pro-inflammatory microenvironment. Cell Death Dis. 2023;14(9):616.

28. Waern I, Lundequist A, Pejler G, Wernersson S. Mast cell chymase modulates IL-33 levels and controls allergic sensitization in dust-mite induced airway inflammation. Mucosal Immunol. 2013;6(5):911–20.

29. Wu N, Zhou T, Carpino G, Baiocchi L, Kyritsi K, Kennedy L, et al. Prolonged administration of a secretin receptor antagonist inhibits biliary senescence and liver fibrosis in Mdr2 -/-mice. Hepatology. 2023;77(6):1849–65.

30. He Z, Song J, Hua J, Yang M, Ma Y, Yu T, et al. Mast cells are essential intermediaries in regulating IL-33/ST2 signaling for an immune network favorable to mucosal healing in experimentally inflamed colons. Cell Death Dis. 2018;9(12):1173.

31. Jalan-Sakrikar N, Anwar A, Yaqoob U, Gan C, Lagnado AB, Wixom AQ, et al. Telomere dysfunction promotes cholangiocyte senescence and biliary fibrosis in primary sclerosing cholangitis. JCI Insight. 2023;8(20).

32. Medina C, Santana A, Paz MC, Diaz-Gonzalez F, Farre E, Salas A, et al. Matrix metalloproteinase-9 modulates intestinal injury in rats with transmural colitis. J Leukoc Biol. 2006;79(5):954–62.

33. Kluger MA, Zahner G, Paust HJ, Schaper M, Magnus T, Panzer U, et al. Leukocyte-derived MMP9 is crucial for the recruitment of proinflammatory macrophages in experimental glomerulonephritis. Kidney Int. 2013;83(5):865–77.

34. Benigni A, Cassis P, Remuzzi G. Angiotensin II revisited: new roles in inflammation, immunology and aging. EMBO Mol Med. 2010;2(7):247–57.

35. Wong CK, Ng SS, Lun SW, Cao J, Lam CW. Signalling mechanisms regulating the activation of human eosinophils by mast-cell-derived chymase: implications for mast cell-eosinophil interaction in allergic inflammation. Immunology. 2009;126(4):579–87.

36. Longley BJ, Tyrrell L, Ma Y, Williams DA, Halaban R, Langley K, et al. Chymase cleavage of stem cell factor yields a bioactive, soluble product. Proc Natl Acad Sci U S A. 1997;94(17):9017–21.

37. Tan N, Lubel J, Kemp W, Roberts S, Majeed A. Current Therapeutics in Primary Sclerosing Cholangitis. J Clin Transl Hepatol. 2023;11(5):1267–81.

38. Goode EC, Rushbrook SM. A review of the medical treatment of primary sclerosing cholangitis in the 21st century. Ther Adv Chronic Dis. 2016;7(1):68–85.

39. O’Brien CB, Senior JR, Arora-Mirchandani R, Batta AK, Salen G. Ursodeoxycholic acid for the treatment of primary sclerosing cholangitis: a 30-month pilot study. Hepatology. 1991;14(5):838–47.

40. Mousavere I, Kalampokis G, Fousekis F, Karayiannis P, Baltayiannis G, Christodoulou D. An overview of recent treatment options for primary sclerosing cholangitis. Ann Gastroenterol. 2023;36(6):589–98.

41. Cure PPSa. Basic PSC Facts: PSC Partners Seeking a Cure; 2025 [Available from: https://pscpartners.org/about/what-is-psc/basic-facts.html.

42. Sankyo D. Study to Evaluate the Efficacy, Safety and Pharmacokinetics of ASB17061 Capsules in Adult Subjects With Atopic Dermatitis clinicaltrials.gov 2014 [Available from: https://clinicaltrials.gov/study/NCT01756898.

